## Supplementary Information, Figures and Tables for "Rapid evolution and biogeographic spread in a colorectal cancer"

**Statistical deconvolution of clonal genotypes from single nucleotide variants**

Clonal deconvolution was performed using the CloneFinder algorithm[1] for the 475 somatic SNVs identified. We required a minimum read count of 40 and a mutant read count of 6 and the clone frequency cutoff was set to 0.075. The 'binary' clonal sequences generated by CloneFinder (i.e., 'A' = reference status; 'T' = alternative alleles) were then modified by changing the binary alleles into the corresponding nucleotide state in the variant allele frequency (VAF) file. We additionally examined the consistency of the clonal genotypes inferred by applying a different clonal deconvolution method, LICHeE[2], to the same dataset. LICHeE was ran using the following thresholds: minVAFPresent = 0.075, maxVAFValid = 0.7, maxVAFAbsent = 0, minClusterSize = 2, minPrivateClusterSize = 1, maxClusterDist = 0.1, outputTrees = 1. The results are illustrated in Supplementary Fig. 3. A total of 18 clones were inferred, displaying a fairly similar phylogenetic relationship to the clones reconstructed using CloneFinder.

**Coding clonal sequences retrieval in PAML**

Codon changes were obtained for each clonal sequence using the *dndscv* R package[3] and concatenated into a multiple sequence alignment. For each branch of the inferred clonal genealogy, we ran PAML 4.8a[4] to obtain maximum likelihood estimates of the non-synonymous/synonymous rate ratio (*dN*/*dS*) using two contrasting models: M0 - which assumes a single *dN/dS* for the whole genealogy, and b_free (i.e., two-ratio) which assumes that the specified branch has a different *dN*/*dS* than the remaining tree branches.

**Spatial distribution of bulk tumor samples**

To assess whether the biogeographical solution described in the main text was robust to changes in the geographical coordinates assigned to each tumor sample, we generated five 2D spatial matrices corresponding to alternative migration distances among the tumor samples (Supplementary Fig. 4). Remarkably, three out of the five 2D matrices resulted in the same migration history as the one described in the main text. Interestingly, for matrix 3, in which the geographical locations of both colonic and hepatic lymph nodes were spaced far apart from the colon and liver, BayArea[5] inferred a biogeographic solution where the ancestral metastatic clone was located in hepatic lymph nodes. In addition, for matrix 5, in which the spatial distance between all organs was substantially reduced, BayArea inferred a migratory dissemination very similar to the one presented in the main text, but suggesting an earlier movement of metastatic clones in the liver to nearby hepatic lymph nodes.

**Inferring migration history using MACHINA**

MACHINA[6] was run in parsimonious migration history mode (i.e., *pmh_sankoff*) using the inferred clonal genealogy from BEAST[7] and setting the colon as the primary anatomical site. As illustrated in Supplementary Fig. 5, MACHINA returned five distinct migratory solutions, displaying a different number of migrations and comigrations. Interestingly, the solutions have several elements in common with the biogeographical scenario inferred using BayArea[5].

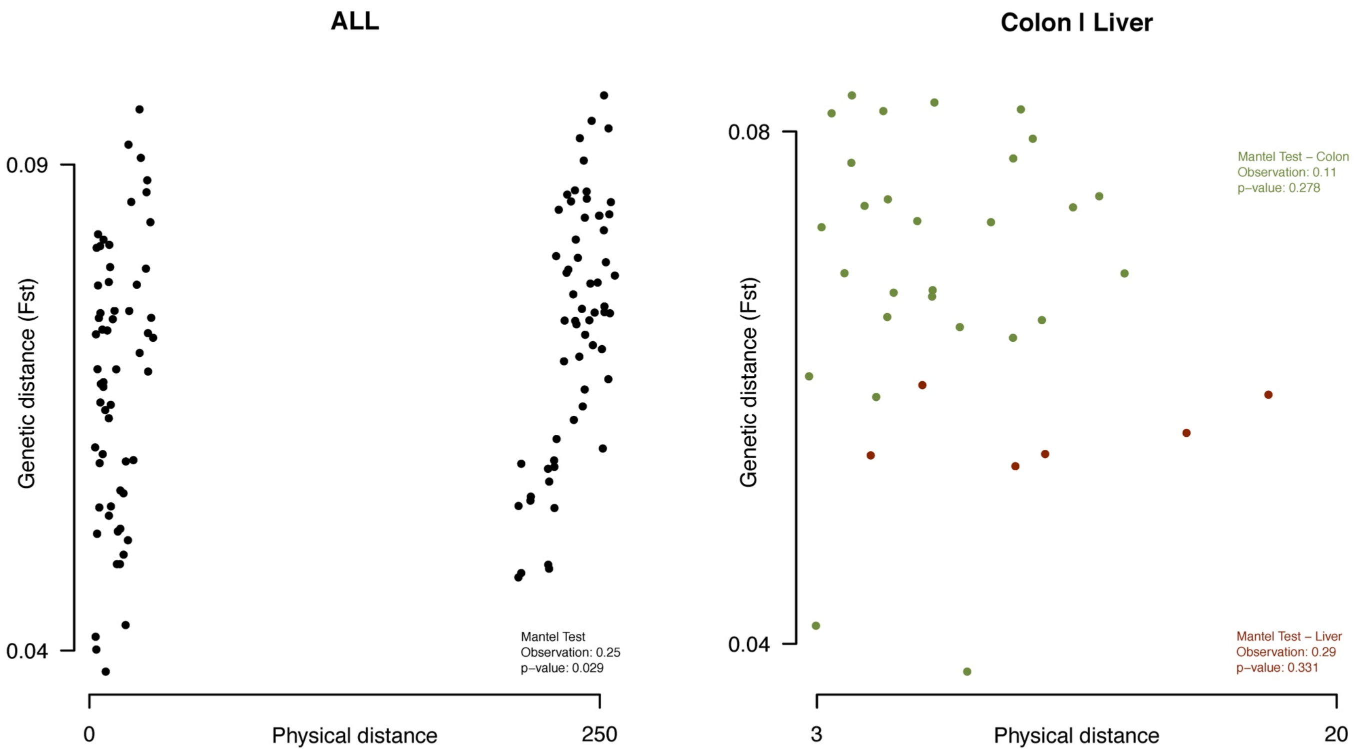


**Supplementary Figure 1. Overall and tissue-specific correlation between geographic distance and genetic distance.** The geographic distance matrix consists of pairwise comparisons of the spatial location of tumor samples in *Matrix 1*. The genetic distance matrix consists of pairwise *Fst* estimates[8]. A Mantel test[9] was performed in R comparing the two distance matrices using 1000 replicates.

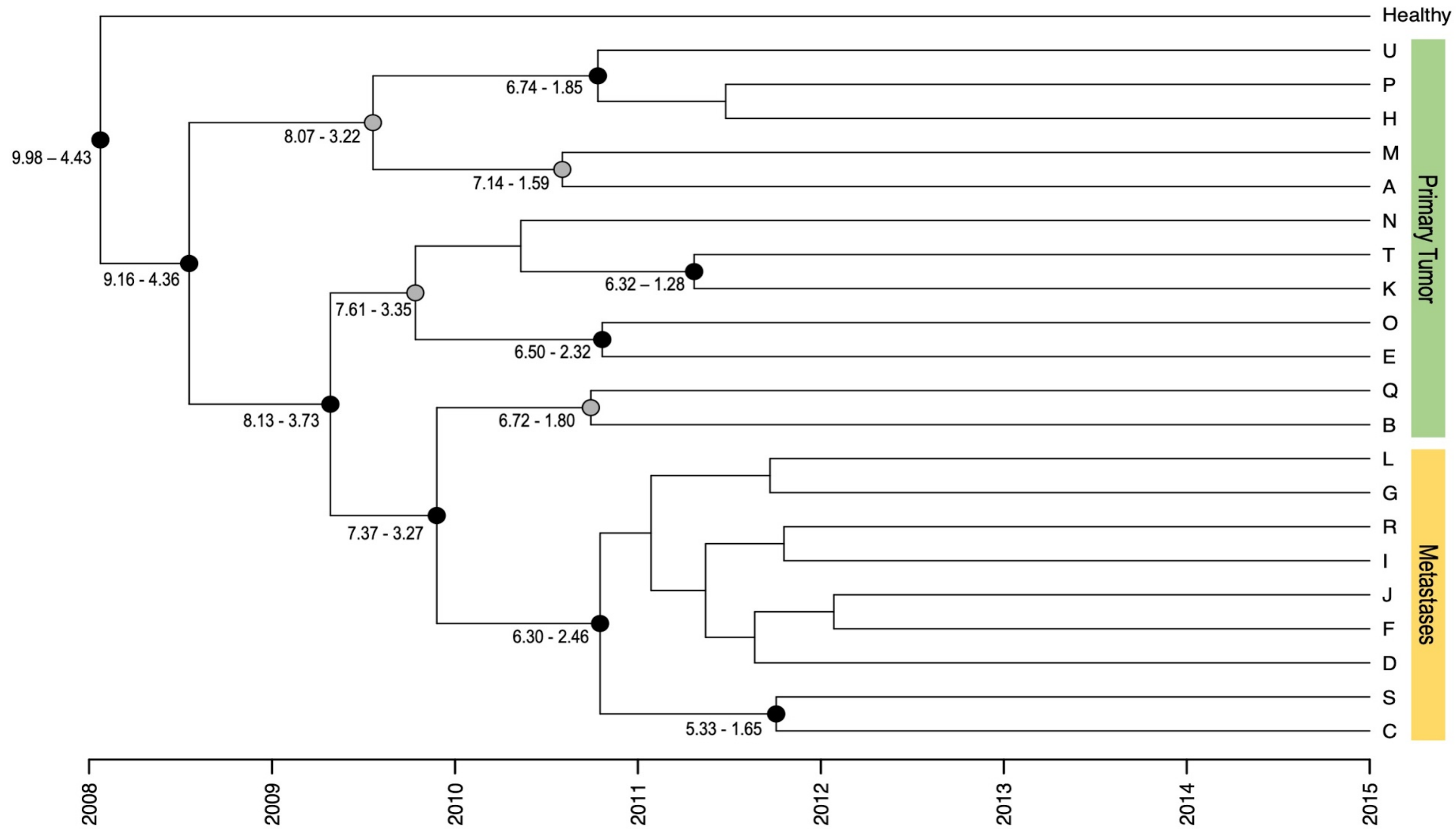


**Supplementary Figure 2. Uncertainty of the phylogenetic dating with *BEAST.** Lower and upper 95% HPD age estimates in years obtained from *BEAST are shown for tree nodes with posterior support > 0.5. Nodes with posterior probability values > 0.9 and > 0.5 are highlighted with black and grey solid circles, respectively. Clone IDs are shown at the tips of the tree.

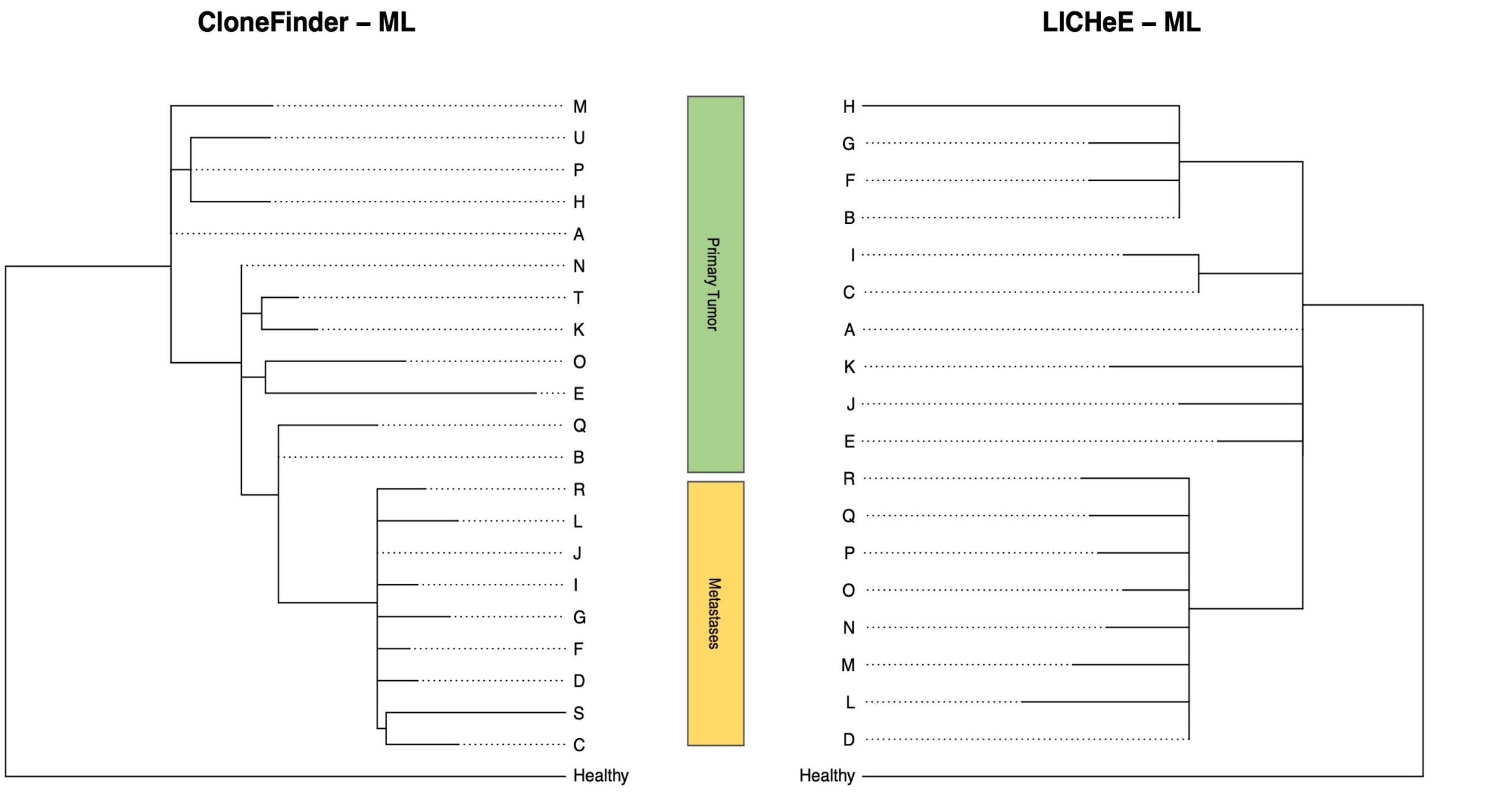


**Supplementary Figure 3. Phylogenetic reconstruction obtained with CloneFinder and LICHeE.** Maximum likelihood trees obtained using heuristic search in PAUP*[10]. Clonal IDs are shown at the tips of the phylogenetic trees (A-U for CloneFinder; A-R for LICHeE). Colored rectangles highlight the anatomical location of each clone: Green - Primary tumor, Yellow - Metastases.

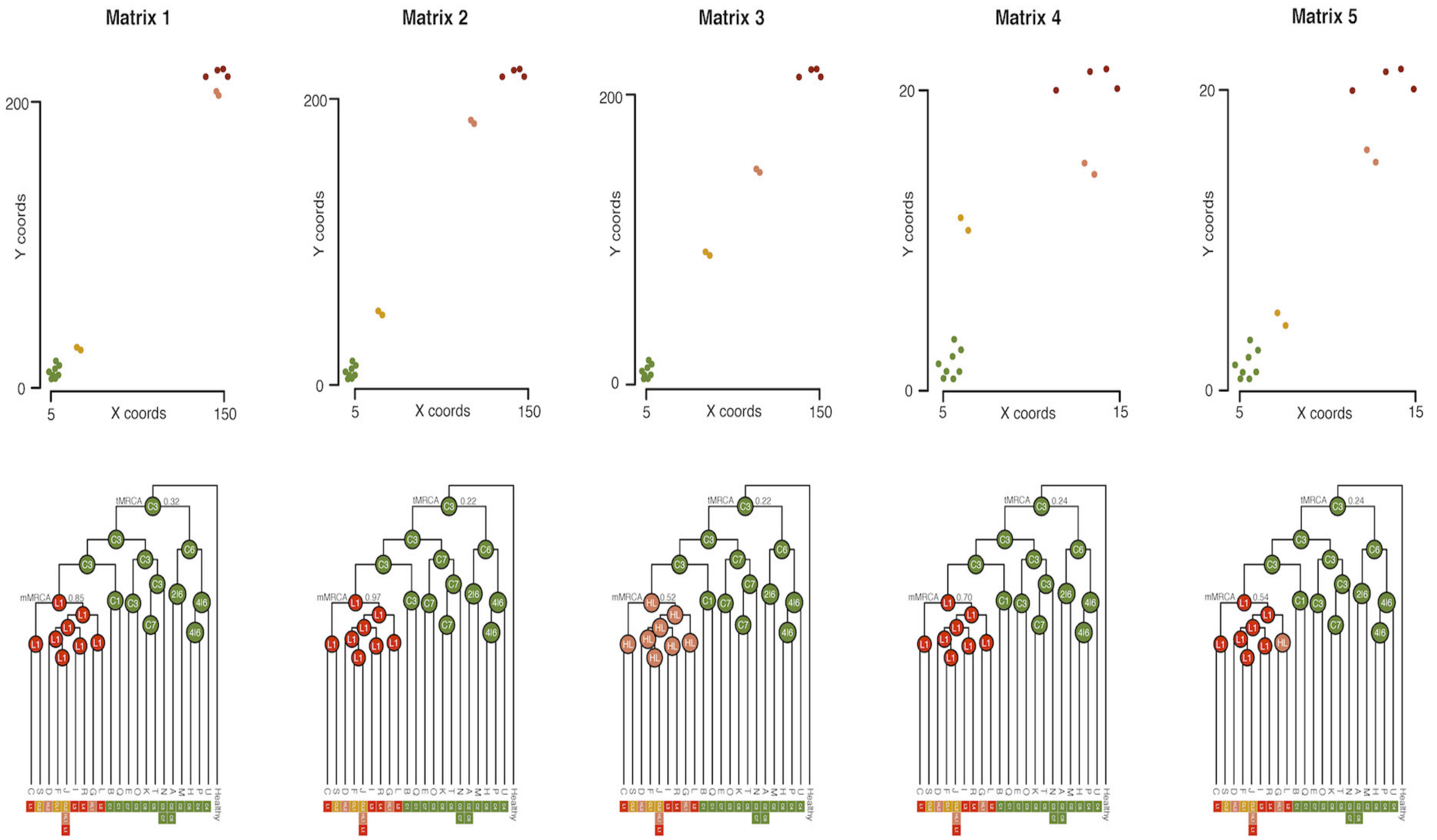


**Supplementary Figure 4. Spatial organization of bulk tumor samples and biogeographic reconstruction. (Top)** 2D coordinate matrices depicting alternative migration tumor samples. Solid circles represent each sample. Colors highlight the anatomical location of each sample: Colon - Green; Colonic Lymph Nodes - Gold; Hepatic Lymph Nodes - Salmon; Liver - Red. **(Bottom)** Biogeographic reconstruction resulting from BayArea using the corresponding 2-D matrix. At two key nodes (tMRCA and mMRCA), the highest posterior probability area range is depicted. Sample IDs are shown at internal nodes. The locations where the extant clones were sampled are shown next to the tips.

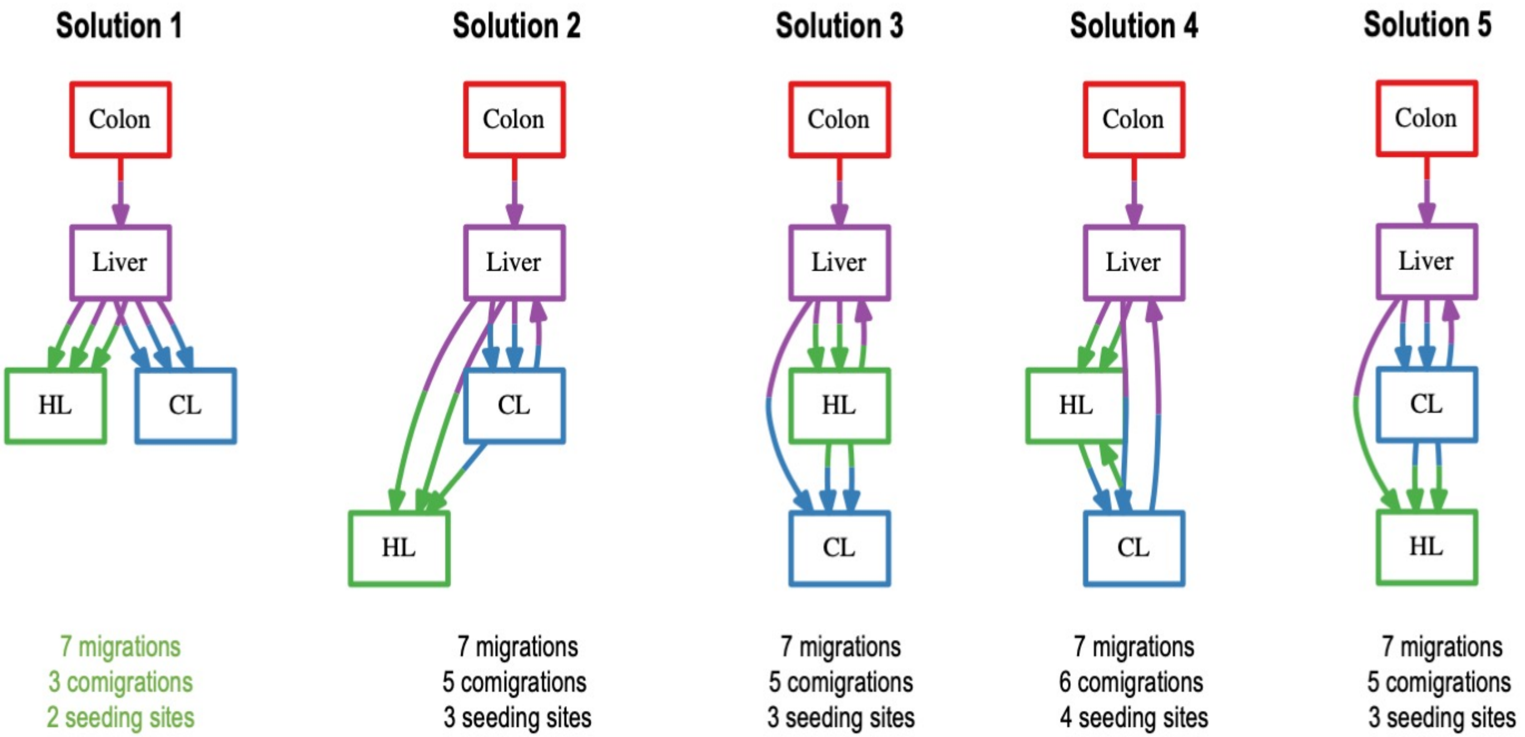


**Supplementary Figure 5. Parsimonious migration inference with MACHINA.** Migration graphs inferred under *phm_sankoff* mode and setting the colon as the primary tumor location. Migratory solutions ordered based on the number of inferred migrations and comigrations. For each graph, colored squares depict the anatomical sites sampled: Colon - red, CL - blue, HL - green, Liver - purple. Arrows indicate clonal movements.

**Supplementary Table 1.** Per-sample variant allele frequency (VAF) for the 475 somatic SNVs.

**Supplementary Table 2.** Evolutionary models tested in BEAST.
